# Menstrual cycle and exogenous attention toward emotional expressions

**DOI:** 10.1101/2022.07.19.500437

**Authors:** Fátima Álvarez, Fernández-Folgueiras Uxía, Constantino Méndez-Bértolo, Dominique Kessel, Luis Carretié

**Affiliations:** Facultad de Psicología, Universidad Autónoma de Madrid, Madrid 28049, Spain

**Keywords:** Exogenous attention, menstrual cycle, emotion, ERP, premenstrual syndrome

## Abstract

Several studies suggest that the menstrual cycle affects emotional processing. However, these results may be biased by including women with premenstrual syndrome (PMS) in the samples. PMS is characterized by negative emotional symptomatology, such as depression and/or anxiety, during the luteal phase. This study aimed to explore the modulation of exogenous attention to emotional facial expressions as a function of the menstrual cycle in women without PMS. For this purpose, 55 women were selected (from an original volunteer sample of 790) according to rigorous exclusion criteria. Happy, angry, and neutral faces were presented as distractors, while both behavioral performance in a perceptual task and event-related potentials (ERPs) were recorded. This task was applied during both phases of the menstrual cycle (luteal and follicular, counterbalanced), and premenstrual symptomatology was monitored daily. Traditional and Bayesian ANOVAs on behavioral data (reaction times and errors in the task) and ERP indices (P1, N170, N2, and LPP amplitudes) confirmed the expected lack of an interaction of phase and emotion. Taken together, these results indicate that women free of PMS present steady exogenous attention levels to emotionally positive and negative stimuli regardless of the menstrual phase.

## Introduction

Automatic or exogenous attention can be understood as an adaptative tool for rapidly detecting salient events and reorienting and enhancing processing resources to them (Carretié, 2014). This cognitive tool is of vital importance for the individual, given that it permits the discrimination and evaluation of whether a stimulus should be urgently attended. Emotional stimuli, which by definition are salient to the individual, are thereby good capturers of exogenous attention. Besides being sensitive to the emotional nature of stimuli, attention capture can be influenced by an individual’s internal predisposition. Thus, exogenous attention toward emotional stimuli may be modulated by affective states or traits such as anxiety, in which negative stimuli capture attention to a greater extent (see a review in Carretié, 2014).

Several studies indirectly suggest that menstrual cycle phases impact women’s mood and affective processing (see Farage et al., 2008 for a review), hence, influencing their exogenous attention to emotional processing. Particularly, an increase of negative affect–such as depressive and anxiety moods-has been described at the end of the cycle or luteal phase (Allen et al., 2009; Ivey & Bardwick, 1968; Li et al., 2020; Reed et al., 2008; Sanders et al., 1983), characterized by an increase of progesterone levels. The hormone fluctuation during the menstrual cycle may underlay women’s regulation of emotion and affect (Andreano & Cahill, 2010; van Wingen et al., 2011). Relatedly, and important to our study, in which emotional faces will be presented to participants, women have been reported to show impairment in facial expression recognition during the luteal phase (see Osório et al., 2018 for a review), suggesting a sex hormone modulation in these tasks (Derntl et al., 2008a; Derntl et al., 2008b; Guapo et al., 2009; Pearson & Lewis, 2005). The increase of physical discomfort and negative emotional symptomatology during the luteal phase is referred to as premenstrual syndrome or PMS (for a review on PMS symptomatology, see Yonkers & Simoni, 2018), whose prevalence is still unclear, affecting from 20 up to 50% of women at their reproductive age (Ryu, 2015; Sattar, 2014; Yonkers, 2018).

Exogenous attention capture may be measured through behavioral and neural indices. At the behavioral level, reaction times and/or number of errors in the ongoing task increase when distractors (irrelevant to the task) capture attention (e.g., Fockert et al., 2004; Hickey et al., 2006; Theeuwes, 1992). At the neural level, event-related potentials (ERPs) are considered a particularly useful technique in the study of rapid emotional and attentional processes thanks to its excellent temporal resolution (Hajcak et al., 2012; Luck et al., 2000). Interestingly, menstrual cycle phases can modulate the amplitude of ERP components indexing attention toward emotional stimuli. For instance, reports exist on a greater amplitude of P1, N1, and N2 (Lusk et al., 2015, 2017; Wu et al., 2014) during the luteal phase towards emotional stimuli. Additionally, N170, a face-sensitive component that has been proven to discriminate among different expressions (see a review in Hinojosa et al., 2015), shows smaller N170 amplitudes to happy expressions during the luteal than during the follicular phase (Yamazaki & Tamura, 2017). Importantly, the majority of these components (namely posterior P1, N2x -or N2 family- and N170), are sensitive to exogenous attention by affective stimuli, showing enhanced amplitudes to emotional distractors -including facial expressions-compared to neutral ones (see reviews in Carretié, 2014; Hinojosa et al., 2015). Therefore, they are appropriate to explore the effect of the menstrual cycle on exogenous attention to emotional faces. Lastly, at later latencies, the Late Positive Potential (LPP) seems to be sensitive to menstrual phase effects -not related to attention capture-showing enhanced amplitudes during the late follicular phase (Krug et al., 2000; Zhang et al., 2013a; 2013b), which can provide us with additional information as an electrophysiological index of hormone-related neural effects.

A question that arises is whether the differences observed in the menstrual-cycle related studies are the consequence of hormone-cycle vulnerability or are the result of negative affective symptomatology of PMS, since women suffering from this syndrome could have been included in the experimental samples. Indeed, the amplitude of P1 and N170 (among those mentioned above) in response to facial expressions are modulated in the general population by mood states such as depression that is often present in PMS (Wu et al., 2016; Zhang et al., 2016). In this respect, several methodological cautions not always followed in previous studies are necessary to explore this issue. First, controlling PMS incidence in the experimental sample seems crucial regarding emotion-related processes (Poromaa & Gingnell, 2014). Thus, it has been recommended to carry out the assessment with a prospective tool during at least two cycles to discard premenstrual disorders (Yonkers & Simoni, 2018). Second, studies on the modulation of emotional processing by the menstrual cycle often employ different samples for each cycle rather than a crossover design, and this could lead to effects related to other inter-individual differences besides hormonal effects. And third, a sample screening to discard medical, psychiatric, and hormone-related disorders, drug consumption (e.g., analgesics or antidepressants), and hormone contraceptive use would also be of interest since they may alter the observed effects.

Our study aimed to characterize the neural and behavioral effects of the menstrual phases on exogenous attention to facial expressions taking these methodological issues into account. For the reasons explained above, we consider that PMS-asymptomatic women may not present differences in emotional processing and, consequently, exogenous attention to emotional stimuli is neither affected. We carried out a within-subject study for three months, monitoring the individual premenstrual symptomatology daily, in order to discard PMS. In our CDTD task *(concurrent but distinct target-distractor task),* optimal for exploring exogenous attention (Carretié, 2014), emotional faces were presented as distractors while participants carried out a perceptual (line orientation) task. ERPs and behavioral performance in the task were recorded. We expected similar reaction times and errors during the follicular and luteal phases for emotional distractors at the behavioral level. Likewise, at the neural level, we expected similar amplitudes in response to emotional distractors in the components mentioned above, reflecting exogenous attention capture: posterior P1 (P1p), N2x, and N170. In other words, we hypothesize no effects of the menstrual cycle on exogenous attention to emotional faces.

## Methods

### 1. Participants

Fifty-five naturally cycling women participated in this experiment, although data from only 43 of them could eventually be analyzed, as explained later (age range of 18-21 years, mean = 19.02, *SD*=0.97; age menarche range of 9-15, mean= 12.11, *SD*= 1.27). The study was conducted according to the Declaration of Helsinki, with approval from the Universidad Autónoma de Madrid’s Ethics Committee. All participants were students of Psychology, provided their informed consent, and received academic compensation for their participation. They reported normal or corrected-to-normal visual acuity.

Potential participants were screened through a self-report questionnaire from a large sample of 790 females. These potential participants were all students at the School of Psychology of the Universidad Autónoma de Madrid receiving a proportional academic compensation, and the questionnaire was designed ad-hoc for sample selection of this study. The selection was carried out according to the following exclusion criteria: non-biological women; either pregnant, nursing, with biological children under one year of age, or who had been pregnant during the year before the study phase; history of neurological, psychiatric, or gynecological diseases; history of hormonal or thyroid-related illness or any disease requiring chronic medication; anxiolytic or antidepressant intake during the last three months; frequent use (more than once in the last month) of other types of medications or recreational drugs; use of hormonal contraceptives in the last three months; irregular average cycle length (less than 24 days or more than 36 days); and women subjectively experiencing PMS as measured through the Premenstrual Symptoms Screening Tool revised for adolescents, PSST-A (Steiner et al., 2011)^1^.

The fifty-five women who survived the exclusion criteria were called to attend an informative meeting about the experiment overview. During two cycles, these women were asked to fill out a Spanish translation of the Daily Record of Severity Problems, DRSP (Endicott et al., 2006), a prospective instrument in order to confirm the non-PMS diagnosis, as a part of a daily self-report. During the informative meeting, the participants were instructed on how to complete the diary (DRSP) through an individual Google Form^®^ (to facilitate accessibility and adherence). Three out of the 55 original participants met the criteria for PMS diagnostic, as revealed by DRSP, despite of having been screened as non-PMS with a previous tool during sample recruit. Due to the diagnosis, these three participants were excluded from further analysis.

### 2. Stimuli and session procedure

Three types of facial expressions were presented to participants: happy (Hap), angry (Ang), and neutral (Neu). These stimuli proceed from the FACES database (Ebner et al., 2010) and depict 50 different young-adult faces (average age of models=24.3, SD=3.5), 25 male and 25 female, all posing the three expressions.

All participants had to attend two experimental sessions: session 1, during the first menstrual cycle (of the two-cycle period we studied), and session 2, during the second cycle. The two dates for the experimental sessions were calculated based on information about the dates and duration of former menstrual cycles. Since these phases depend on the menstrual cycle length of the participants (mean= 28.53, *SD*=1.86), the measurement moment was adjusted individually. For the follicular phase (Fol), participants were appointed between day 6 to 12 after the beginning of menses (mean= 9.72, *SD*= 2.12). For the luteal phase (Lut), they were appointed 7-2 days before the expected start of the next menses, which, depending on each participant’s cycle length, corresponded to days 20-28 of their cycle (mean= 24, *SD*= 2.15). Retrospectively, the actual day of each cycle was confirmed based on the daily self-reports. Two participants reported that their menses had started, some hours later, on the same day their Lut session took place, so both were discarded for further analyses according to the criteria proposed by Schmalenberger et al. (2021). For the rest of participants, the Lut session actually took place between the previous 12 days and the day before the beginning of the next cycle (mean=5.16, *SD*=2.74). We also confirmed based on the self-reports that participants in Lut session were in the last third of the cycle in progress, therefore, it was highly probable that ovulation had already taken place. The order of Lut and Fol for sessions 1 and 2 was counterbalanced^2^. The mean interval between EEG sessions 1 and 2 was 43.96 days (*SD*= 8.49).

In each of the two sessions, participants were placed in an electromagnetically shielded, sound-attenuated room. They were asked to sit on a chair, maintained at a fixed distance of 85 cm from the screen (VIEWpixx^®^, 120 Hz). Faces were presented as distractors in the center of the screen, adjusted to a width of 10.422° and height of 13.022°, and presented on a black background. We programmed our experimental task in Matlab using Psychtoolbox extensions (Brainard, 1997). Targets consisted of two yellow lines (width 0.67° and height 11.85°) presented on both sides of each face and showing variable orientations. The orientation of both lines was either equal or different (50%-50%; in the latter case, whatever their orientation, the difference was always 36°). The total set of 300 trials (100 per emotion) was displayed randomly and separated into two blocks of 150 trials to provide a brief rest. Participants were instructed to indicate, in each trial, whether the two lines had the same or different orientations by pressing two different keys on a numbered keyboard (the key assignment was counterbalanced across participants). A practice block of 10 trials presenting different faces to those presented in the experimental run was previously administered to ensure that the instructions had been understood. Each stimulus was displayed for 300 ms, and the intertrial interval (in which a yellow fixation dot -0.30° diameter- on a black background was displayed) lasted randomly between 1350 and 1850 ms (Figure 1). Participants were instructed to permanently direct their gaze to the fixation dot and to avoid any ocular movement. The total duration of the stimulus sequence was between 8.25 to 10.75 minutes. Two out of the 55 original participants did not attend their second session and were not analyzed.

**Figure 1.**
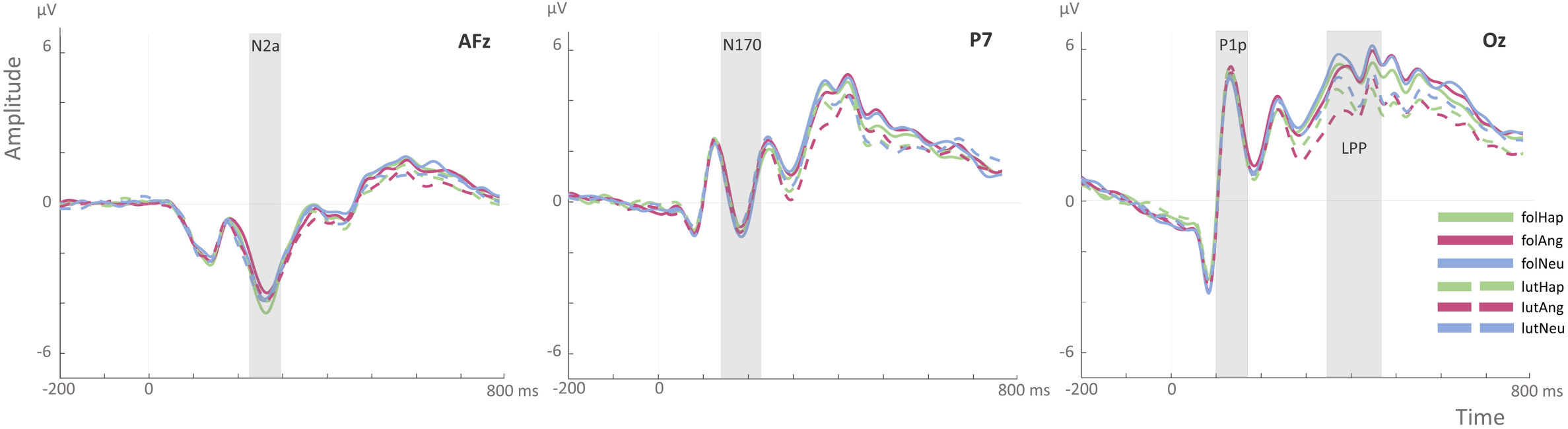
Schematic illustration of the CDTD task showing the first trial, after 250 ms of the blank. The subsequent trials were displayed after a randomly intertrial interval between 1350 and 1850 ms. Notice that the distractor in the first example trial is an angry face with similar line orientation, while the following trial displays a happy face as a distractor with different line orientations.

### 3. EEG recording, preprocessing, and control analyses

Electroencephalographic (EEG) activity and behavioral data were collected during the two sessions. The task response was collected through a numbered keyboard. EEG activity was recorded using an electrode cap (ElectroCap International) with tin electrodes. Fifty-nine electrodes were placed at the scalp following a homogeneous distribution and the International 10-20 System. All scalp electrodes were referenced online to the nose tip^3^. Supra- and infraorbital electrooculography (vertical EOG) data were recorded, as well as from the left versus right orbital rim (horizontal EOG) to detect blinking and ocular deviations from the fixation point. An online analog high-pass filter was set to 0.03 Hz. Recordings were continuously digitized at a sampling rate of 420 Hz. An offline digital Butterworth bandpass filter (order: 4, direction: zero-phase forward and reverse, two pass-filter) of 0.3-30 Hz was applied to continuous (preepoched) data using the Fieldtrip software (Oostenveld et al., 2011). The continuous recording was divided into 1000 ms epochs for each trial, beginning 200 ms before the stimulus onset.

Ocular artifact removal was carried out through an Independent Component Analysis (ICA)-based strategy (Jung et al., 2000), as provided by Fieldtrip. After the removal process, a visual inspection of EEG data was conducted in order to manually discard trials in which any further artifacts were present. The minimum number of trials accepted per participant and condition was 51 (50%). Five out of the participants excluded for analyses did not reach this threshold or presented data loss. This automatic and manual rejection procedure led to an average admission of 83.8 trials (*S*D= 9.21) in FolHap trials, 84.8 (*SD*=7.82) in FolAng, 84.3 (*SD*=7.91) in FolNeu, 81.5 (*SD*=11) in LutHap, 80.3 (*SD*=11.2) in LutAng, and 80.0 (*SD*=11.5) in LutNeu trials. The difference of accepted trials among the six conditions was non-significant (F (2, 84) =2.58, Greenhouse-Geisser corrected *p*= 0.08, η^2^□= 0.06). Besides these correction and rejection strategies, additional control analyses were carried out in order to discard any significant influence of ocular activity in the observed results. To that aim, original data from both horizontal and vertical EOG channels (hEOG and vEOG) were submitted to Friedman’s non-parametric test, since they did not achieve normality (S-W tests of normality, *p* < 0.001 in both cases). The differences regarding ocular movements among conditions were not significant, neither the hEOG (Friedman’s test: χ^2^(5) =2.53, *p*=0.771) nor the vEOG (Friedman’s test: χ^2^(5) =2.35, *p*=0.799).

### 4 Data analysis

#### 4.1 Detection, spatiotemporal characterization, and quantification of relevant ERP components

Detection and quantification of the ERP components taken into account in our hypotheses were carried out via two-step covariance-matrix-based principal component analysis (PCA): a temporal PCA (tPCA) followed by a spatial PCA (sPCA)^4^. PCA has repeatedly been recommended for data reduction purposes to differentiate individual ERP components and to handle component overlap (e.g., Chapman et al., 2004; Dien, 2010). This technique defines temporal windows and spatial regions mathematically, based on the covariance of amplitudes both in time and in space, avoiding subjectivity or inter-judge discrepancies often characterizing the traditional window and region definition based on manual or visual criteria. Thus, it guarantees more objective and reliable results than traditional, visual inspection-based methods. In both steps, the decision on the number of factors to select in the two PCAs was based on the scree test (Cliff, 1987), and extracted factors were submitted to Promax rotation also in both cases (Dien, 2010). This analysis was performed using SPSS 26.0 software package (IBM Corp., 2019).

Data reduction in the time domain: tPCA. Firstly, detection and quantification of prominent perception-related temporal components were carried out through a covariancematrix-based temporal PCA (tPCA). In brief, tPCA computes the covariance between all ERP time points (each 1000 ms epoch comprises 420 time points, as previously explained) across participants and conditions, which tends to be high between those involved in the same component and low between those belonging to different components. The solution is therefore a set of nearly independent factors made up of highly covarying time points, which ideally correspond to ERP components. The matrix submitted to tPCA was formed by voltages as variables and participants × emotion × phase × channels (i.e., 45 × 3 × 2 × 59 = 15930) as cases. Among the resulting temporal factors (TFs) selected through the scree test previously mentioned, those whose latency was equivalent to the components mentioned in the Introduction, were selected. Extracted temporal factors (TF) were quantified as factor scores, which are linearly related to amplitudes: original amplitudes are a direct function of factor loadings (the global weight of each data point within each TF) and individual factor scores (Dien, Tucker, Potts, & Hartry-Speiser, 1997; Dien et al., 2005, 2007). The TF scores, consisting of a single value per TF, were submitted to the next PCA step.

Data reduction in the topography domain: sPCA. Secondly, TF5, TF6, TF4, and TF2 factor scores resulting from the previous tPCA were submitted to sPCA in order to decompose topographies at the scalp level into their main spatial regions. Thus, while tPCA separates ERP components along time, sPCA reliably separates them in space, each region or spatial factor (SF) ideally reflecting one of the concurrent neural processes underling each temporal factor. This spatial PCA provides a reliable division of the scalp into the different regions or spatial factors (SFs) in which each TF is distributed. Basically, each SFs is formed by the scalp points where recordings tend to covary. So, the shape of the sPCA-configures regions is functionally based. Similarly, the decision on the number of factors to select was based on the scree test and extracted factors were submitted to promax rotation as well. Each input matrix (one for each of the four selected TFs) consisted of 59 variables (i.e., EEG channels) and 270 cases (i.e., participants × emotion × phase). Statistical analyses were computed on SF scores, which are linearly related to amplitudes.

#### 4.2 Analyses of experimental effects

Regarding the behavioral data, errors -defined as incorrect responses- and reaction times (RTs), measured in seconds, were analyzed. Before that, trials in which RT were three standard deviations below or above the individual RT mean were discarded. Both behavioral measures were submitted to non-parametric contrasts, due to their non-Gaussian distributions (K-S test on number of errors D (258) = 0.113, *p* < 0.001; K-S test on reaction times: D (258) = 0.106, *p* < 0.001). In order to test possible interactions, which was our main concern, Fol minus Lut subtractions were computed both for errors and RTs: FolHap-LutHap, FolAng-LutAng, FolNeu-LutNeu. These three levels were submitted to a Friedman’s test. Additionally, main effects were also tested by submitting Emotion (Hap, Ang, Neu) and Phase (Fol, Lut) to Friedman’s tests on error and RTs. In case of significance of the Friedman’s test, post hoc pairwise comparisons were computed employing the Wilcoxon signed-rank test. These analyses were carried out using SPSS 26.0 software (IBM Corp., 2019).

Concerning ERP analyses, experimental effects on relevant components were tested by introducing Phase (Fol, Lut) and Emotion (Hap, Ang, Neu) as within-subject factors in a repeated-measures ANOVAs performed on spatial factor scores, corresponding to amplitudes of each ERP component at each topography, as above-mentioned, using AFEX R (Singmann, 2018) and Jamovi R Statistical Software packages (The jamovi proyect, 2021; R Core Team, 2020). We used the Greenhouse-Geisser (G-G) epsilon correction to adjust the degrees of freedom of the F ratios if necessary, and post hoc comparisons were performed to determine the significance of pairwise contrasts applying the Bonferroni correction procedure. Effect sizes were computed using the partial eta-square (η^2^_p_) method.

Additionally, Bayesian Repeated Measures ANOVAs computed via JASP (JASP Team, 2020) were carried out to obtain evidence in favor of the null hypothesis (H0: Fol=Lut for any level of Emotion) over the alternative hypothesis (H1: Fol ≠ Lut in some level of Emotion) in the interaction analysis between phase and emotion. The Cauchy prior values were set to 0.5 for both behavioral measures (errors and RT), expecting a medium effect size, in concordance with previous studies (Pearson & Lewis, 2005; Zhang et al., 2016). Regarding the neural data, the priors were set to 1 according to the effects sizes reported in previous studies on the LPP component (Lustig et al., 2018; Zhang et al., 2015) and N2x (Lusk et al., 2015; Wu et al., 2014; Zhang et al., 2015), and set to 0.5, conforming to the effects sizes previously reported on N170 (Hinojosa et al., 2015; Lustig et al., 2018; Zhang et al., 2016), and P1 (Lusk et al., 2015, 2017; Zhang et al., 2016). A Bayesian Factor (BF_10_) over 3 represents evidence supporting H1 over H0, and less than 1/3 evidence in the opposite direction: H0 over H1 (Jeffreys, 1939; see also Dienes, 2014).

## Results

### 1. Experimental effects on behavior

Means and standard deviations of behavioral data are presented in Table 1. Results regarding errors revealed a non-significant interaction between Phase and Emotion computed via the Fol-Lut subtractions described in the previous section (Friedman’s test: χ^2^= 5.35; *p* = 0.069). In addition, Bayesian ANOVAs regarding the phase and emotion interaction found substantial evidence in favor of the null hypothesis (BF_10_=0.136). No main effect of Emotion (Friedman’s test: χ^2^= 1.90 *p* = 0.387) was found. However, a main effect of Phase reached significance (Friedman’s test: χ^2^= 3.90, *p* = 0.048), where the Fol condition (mean= 6.15, *SD*=4.27) caused a greater number of errors than Lut (mean= 5.69, *SD*=4.25).

**Table 1.** Means and standard deviations (in parenthesis) of (i) number of errors and (ii) reaction times.

|  | FolHap | FolAng | FolNeu | LutHap | LutAng | LutNeu |
| --- | --- | --- | --- | --- | --- | --- |
| Errors | 6.09 (3.96) | 6.00 (4.39) | 6.34 (4.55) | 5.20 (4.25) | 6.00 (4.40) | 5.86 (4.16) |
| Reaction times<br>(seconds) | 0.691<br>(0.115) | 0.695<br>(0.121) | 0.688<br>(0.118) | 0.715<br>(0.130) | 0.718<br>(0.132) | 0.714<br>(0.134) |

RT did not show any significant interaction effect (Friedman’s test: χ^2^=0.884, p=0.643), and Bayesian analysis confirmed strong evidence in favor of the null hypothesis (BF_10_=0.078) regarding the interaction. No main effect of Emotion was found (Friedman’s test: χ^2^=2.23, p=0.327), nor a main effect of Phase (Friedman’s test: χ^2^=3.42, p=0.064).

### 2. ERPs: identification and characterization of relevant ERP components

Figure 2 shows a selection of grand averages from each ERP. These grand averages correspond to anterior (AFz), left parietal (P7), and posterior (Oz) channel, where relevant components were identified. As previously indicated, the first analytical step consisted of detecting and quantifying the relevant components through tPCA (see the section on Data Analysis). As a consequence, six temporal factors (TF) were extracted by tPCA and submitted to Promax rotation. As Figure 3 shows, four of them corresponded to the components of interest (mentioned in the introduction), based on their factor peak latency. Thus, TF5, TF6, TF4, and TF2 were associated with latencies of P1p (peak latency =≃ 130 ms), N170 (peak latency =≃ 170 ms), N2x (peak latency ≃ 250 ms), and LPP (peak latency ≃ 470 ms) and considered for further analyses, as indicated. The second step, as also indicated, consisted of applying sPCAs to each TFs to determine their spatial distribution. With this aim, TF5 (P1), TF4 (N2), and TF2 (LPP) were decomposed into two SFs, one anterior and one posterior in all cases, and TF6 was decomposed into five SFs that included those corresponding to the typical left and right parieto-temporal distribution of N170 (Figure 3).

**Figure 2.**
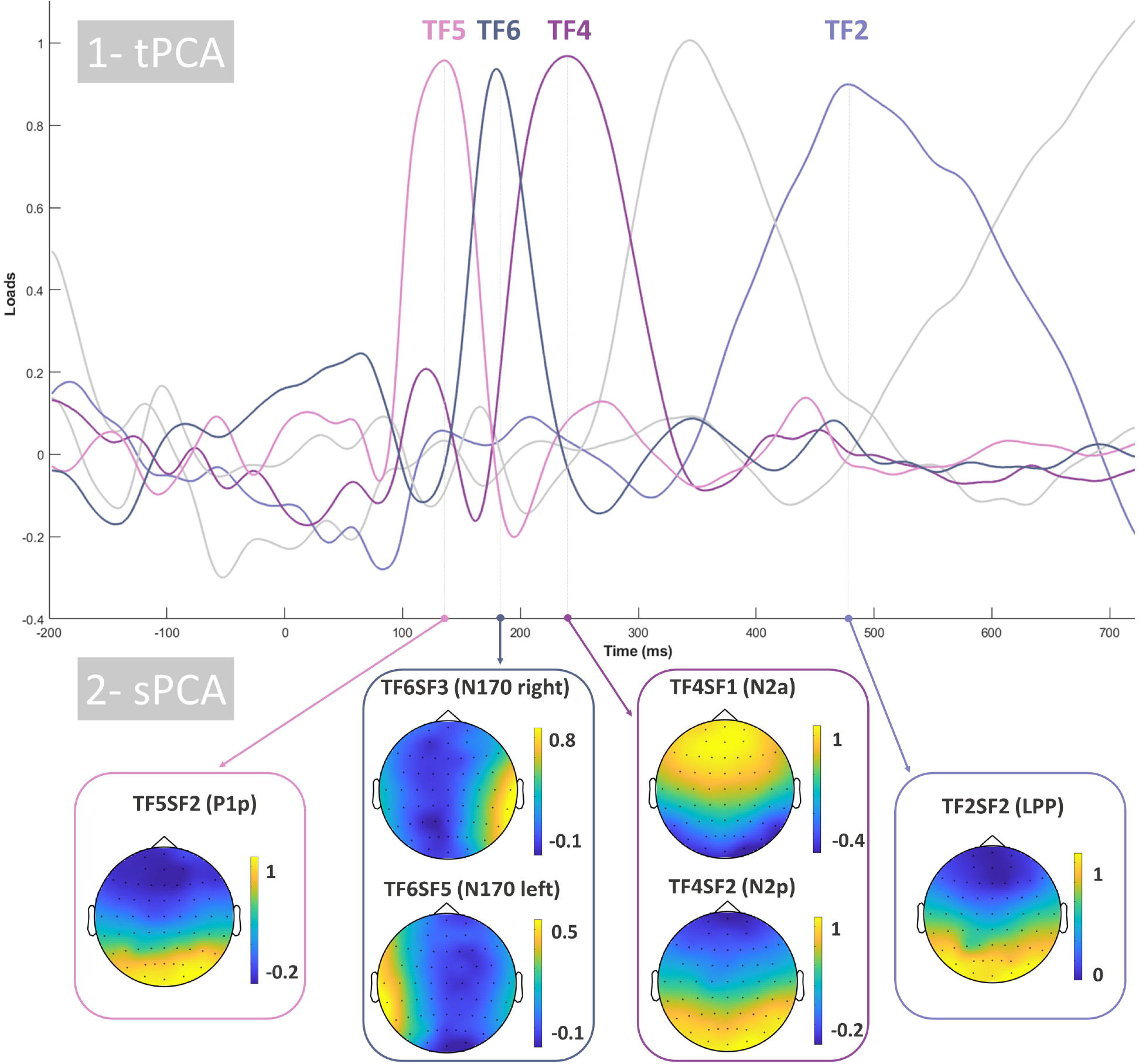
Grand averages corresponding to anterior (AFz), left parietal (P7), and posterior (Oz) electrodes for the components which were previously identified: P1p, N170, N2a, and LPP.

**Figure 3.** Two-step principal component analysis (PCA) structure. First, temporal PCA (tPCA) extracted temporal factors (TF) from original recordings, being P1 (TF5), N170 (TF6), N2 (TF4), and LPP (TF2) those relevant to this study (highlighted in color). The second step consisted of submitting the TF scores to spatial PCAs (sPCA) to decompose them into spatial factors (SFs). Only relevant spatial factors are shown, see main text for details. Please note that both temporal and spatial factors are positive loads.

### 3. Experimental effects on ERP components

Table 2 shows the means and standard deviations of the main SF scores (equivalent to amplitudes, as indicated) corresponding to P1p, N170, N2x, and LPP in each experimental condition. These factor scores were submitted to repeated-measures ANOVAs (as explained in the Methods section).^5^

**Table 2.**
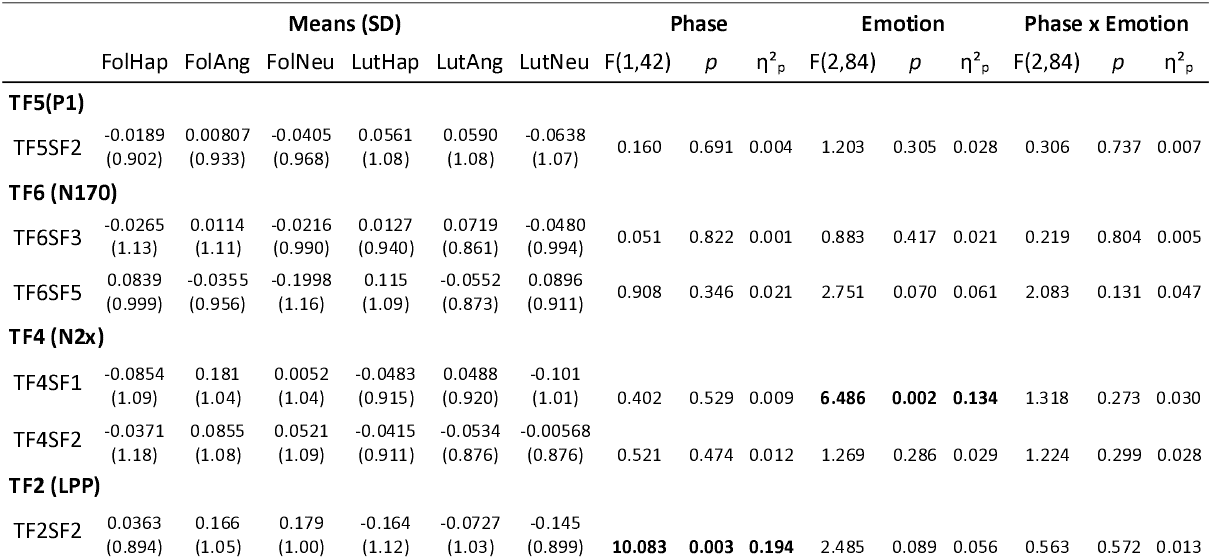
Means and standard deviations (in parenthesis) of SF scores for P1p, N170, N2x, and LPP (linearly related with amplitudes) corresponding to each experimental condition. Results of the two-way ANOVA with Phase and Emotion as factors. p values were GG-corrected when necessary. Significant and relevant results are shown in bold.

#### P1p (TF5; peak at 130 ms)

The relevant spatial factor was SF2, which showed a posterior topography. Results on this SF did not reveal any significant Phase × Emotion interaction [F (2, 84) = 0.306, *p* = 0.737, η^2^_p_ = 0.007], main effect of Phase [F (1, 42) = 0.160, *p* = 0.691, η^2^_p_ = 0.004], nor Emotion [F (2, 84) = 1.20, *p* = 0.305, η^2^_p_ = 0.028]. Bayesian ANOVAs regarding the interaction effect also confirmed strong evidence in favor of the null hypothesis (BF_10_=0.0973). Consequently, this component was discarded for further analyses.

#### N170 (TF6, peak at 170 ms)

Relevant spatial factors were SF3 and SF5, which corresponded with the widely described topography of N170: right and left temporo-parietal distributions, respectively. Results on SF3 did not reveal any significant Phase × Emotion interaction [F (2, 84) = 0.219, *p* = 0.804, η^2^_p_ = 0.005], main effect of Phase [F (1, 42) = 0.051, *p* = 0.822, η^2^_p_ = 0.001], nor Emotion [F (2, 84) = 0.883, *p* = 0.417, η^2^_p_ =0.021], Bayesian repeated measures ANOVA confirmed strong evidence in favor of the null hypothesis regarding the interaction effect (BF_10_=0.0896). The left temporal distribution of N170, reflected in SF4, either showed any significant effect of Phase × Emotion interaction [F (2, 84) = 2.08, *p* = 0.131, η^2^_p_ = 0.047]. Bayesian ANOVAs regarding the phase and emotion interaction found substantial evidence in favor of the null hypothesis (BF_10_=0.303). No main effect of Phase [F (1,42) = 0.908, *p* = 0.346, η^2^_p_ = 0.021], nor Emotion [F (2, 84) = 2.75, *p* = 0.07, η^2^_p_ = 0.061] was found. Therefore, this component was not subjected to further analysis.

#### N2x (TF4, peak at 250 ms)

This component was decomposed into two SFs. On the one hand, SF1, from this point forward N2a, showed an anterior topography. Data Analysis revealed no Phase × Emotion interaction [F (2, 84) = 1.318, *p* = 0.272, η^2^_p_ =0.030] in SF1. Bayesian analysis regarding the phase and emotion interaction confirmed strong evidence in favor of the null hypothesis (BF_10_=0.0447). No main effect of Phase [F (1, 42) = 0.402, *p* = 0.529, η^2^_p_ =0.009] was found. However, a significant effect of Emotion was observed [F (2, 84) = 6.486, *p* = 0.002, η^2^_p_ =0.134], Post-hoc comparisons showed significant differences consisting in Hap>Ang (t (84) = −3.309, Bonferroni corrected p=0.006) and Neu>Ang (t (84) = 2.788, Bonferroni corrected p=0.024) where greater amplitudes consisting of more negative values in this negative component. On the other hand, the ANOVA on SF2 (occipital distribution) did not reveal any significant main or interaction effect (see details in Table 2). The Bayes Factor also confirmed strong evidence in favor of the null hypothesis regarding the interaction effect (BF_10_=0.0375).

#### LPP (TF2, peak at 470 ms)

The relevant SF, characterized by a posterior topography typically associated with LPP, was SF2. This factor showed no Phase × Emotion interaction [F (2, 84) = 0.563, *p* = 0.572, η^2^_p_ = 0.013]. In addition, Bayesian analysis of the interaction between phase and emotion showed strong evidence in favor of the null hypothesis (BF_10_=0.0326). A significant main effect of Phase [F (1, 42) = 10.083, *p* =0.003, η^2^_p_ = 0.194], but not of Emotion [F (2, 84) = 2.485, *p* = 0.089, η^2^_p_ = 0.056], was found. Post-hoc comparisons showed significantly greater LPP amplitudes in Fol than Lut (t (42) = 3.18, Bonferroni corrected p=0.003).

## Discussion

This study explored whether exogenous attention to emotional stimuli (facial expressions) is modulated by the menstrual cycle in women after discarding premenstrual syndrome. At the behavioral level, we expected no differences in reaction times and errors in the exogenous attention task toward facial expressions as a function of the menstrual phase, which should mean a lack of menstrual cycle effects, at least in asymptomatic women. Similarly, at the neural level, ERP indices of exogenous attention were expected not to differ as a function of the menstrual phase. Importantly, a PMS diagnosis was discarded through a prospective two-month diary (DRSP, Endicott et al., 2006) for all the analyzed participants, and the results were supported by additional analysis using different reference electrodes and analytical procedures (<u>Supplementary Material</u>). Our results show the absence of menstrual phase modulation of exogenous attention to emotional distractors when PMS is controlled in the behavioral and the neural domain, confirming our hypothesis.

Regarding the behavioral results, we found a main effect of the menstrual phase in the number of errors during the task. Concretely, the follicular phase was associated with more errors than the luteal phase. Importantly to our scopes, this effect did not interact with the emotional content of distractors. Sex hormones are being related to impulsivity and risk-taking (for a review, see Kurath & Mata, 2018), higher estradiol levels being associated with less response inhibition (Colzato & Hertsig, 2010; Protopopescu et al., 2005), which could lead to shorter reaction times and the increase of errors during the task. In fact, behavioral effects have been steadily reported in the follicular phase (Kumar et al., 2013; Souza et al., 2012; Yamazaki & Tamura, 2017), and, important to our study, even when PMS is discarded (Hoyer et al., 2013). Therefore, this result could be taken as a faint evidence to support participants were in their follicular phase as they indicated through of the diary report.

Concerning the neural data, we found lack of modulation of exogenous attention to emotional stimuli by the menstrual cycle. Importantly, Bayesian analyses confirmed the null effect, supporting our hypotheses. These results were also confirmed by the traditional analysis on the common average reference and the nose tip reference (see results in <u>Supplementary Material</u>), pointing to a consistent finding even employing pre-processing and analytical procedures more prone to lead to significant differences in statistical contrasts. These findings open the question on whether previous results regarding emotional processing and the menstrual cycle could be influenced by the inclusion of women with PMS in the sample and highlights the need to control for PMS, in line with our hypothesis. The present data confirm that attention to emotional stimuli is not influenced by the phase of the menstrual cycle, at least at the exogenous level and when stimuli are faces. Further, secondary results are worth to be discussed. Thus, we found an emotion effect at early latencies (N2a) and a phase effect at late latencies (LPP), both components hence reflecting independent processes, each one underlying one of the factors manipulated in this study. Firstly, a main effect of emotion was found in the N2 family of components, specifically, in the N2a. Whereas this anterior distribution it is not the most commonly reported, the results are in line with previous studies that describe N2a modulations as a function of the emotional content of faces (e.g., Kiss & Eimer, 2008; Zhang, 2013b, 2015) and, particularly in line with studies exploring exogenous attention to emotional scenes (Carboni et al., 2017; Kosonogov et al., 2019) or facial expressions (Balconi & Pozzoli, 2008; Streit et al., 2001; Wynn et al., 2008). Therefore, N2a results confirmed that non-negative distractors (concretely, neutral and positive faces) captured attention to a greater extent than negative in our CDTD task, in line with previous studies reporting enhanced N2a amplitudes toward positive distractors (Carboni et al., 2017). This result could be due to the positivity offset effect (Cacioppo & Gardner, 1999), a processing bias favoring the processing of positive stimuli. This processing bias appears in low arousing situations (e.g., when stimuli or context are not emotionally intense). Eventually, we did not find any phase or phase x emotion interaction effect in this component. In this respect, the sparse literature about the influence of menstrual cycle in N2x (not specifically anterior but with diverse topographies) in response to emotional visual stimuli provides mixed results: several studies, in line with ours, have failed to find a menstrual cycle modulation of N2x (Kranczioch et al., 2016; Lusk et al., 2015, 2017; Zhang et al., 2015; 2013b), while one other reports larger N250 amplitudes to moderately and highly negative stimuli during the luteal phase (Wu et al., 2014). However, some differences regarding the methods employed in this study could also explain the results: firstly, the emotional stimuli in the cited study were attended during the task, thus, leading to conclusions at the endogenous attentional level; secondly and importantly, it did not screen the sample for PMS, thus, the alternative interpretation of PMS explaining the results on N250 cannot be ruled out.

Moreover, the LPP was modulated by the menstrual phase of participants, presenting higher amplitudes during the follicular phase. This result is in line with previous studies that have reported higher LPP amplitudes during the periovulatory phase (Krug et al., 2000; Zhang et al., 2013b), which lay within a late follicular phase in terms of hormone levels, similarly to our study, when participants were tested days 6 to 12 of the cycle. In addition, a correlation between the amplitude of the LPP and ovarian hormone levels has previously been reported (Mačiukaitė et al., 2017; Munk et al., 2018; Zhang et al., 2015). However, we need to interpret this result with caution, given that this would be only an indirect measure that may not replace direct, hormonal measures. Additionally, this LPP effect was confirmed by traditional analyses on nose tip reference data, but not by those on common average reference data (<u>Supplementary Material</u>). In any case, and importantly, we did not find any menstrual cycle modulation effect on the LPP amplitude in response to facial expressions, in line with previous studies that explored this issue (Lustig et al., 2018; Zhang et al., 2015; 2013b) and as confirmed by alternative analyses described in <u>Supplementary Material</u>.

In sum, this study finds no modulation of exogenous attention to emotional faces by the menstrual cycle, in line with our hypothesis. Some limitations should, however, be considered when interpreting the results and in future research in this field. Firstly, the lack of hormone analysis deprived us of an objective measure of the phase of the menstrual cycle, which was mainly based on participants’ self-reports. While our study certainly does not focus on specific phases of the menstrual cycle (e.g., the periovulatory phase, for which hormone analysis would be essential), but rather on the comparison of very long phases (whose beginning and the end are well defined by ovulation and menstruation), objective hormonal measures would have benefited our research and would be of great interest in future studies. Secondly, this initial step to explore the effect of the menstrual cycle controlling for confounding effects of PMS opted for excluding this syndrome from our experimental sample. This strategy has been efficient by revealing that the menstrual cycle per se does not inherently affect emotional processing (at least at the attentional capture level). However, exploring the effect of both luteal and follicular phases in attentional capture by emotional distractors in women presenting PMS would be a valuable forthcoming step within this field despite the potential risks that a between-subject design could imply (such as individual differences regarding other potential interfering factors).

## Conclusion

In summary, our study shows the absence of menstrual phase modulation of exogenous attention to emotional distractors in the neural domain in non-PMS women, at least when the distractor stimuli consist of facial expressions. To the best of our knowledge, this is the first study exploring exogenous attention modulation by the menstrual cycle toward expressions, which provides a novel line of research regarding women’s health and wellbeing. Furthermore, our study highlights the importance of controlling for a possible effect of premenstrual syndrome, and the importance of confirming the diagnosis through a prospective tool, when investigating the influence of the menstrual cycle on emotion processing.

## Acknowledgments

This research was supported by the Ministerio de Ciencia, Innovación y Universidades (Grant no. PGC2018-093570-B-100) and the Comunidad de Madrid (Grant no. HUM19-HUM5705) in collaboration with the Universidad Autónoma de Madrid (Grants no. 2017-T2/SOC-5569 and S|1-PJ|-2019-00011).

## Footnotes

1 PSST-A items are the same as those of PSST (Steiner, 2003), except the former checks for the interference of PMS symptoms in the academic context instead of the professional context. Given that our sample consisted of students, the former was considered more convenient.

2 A saliva sample was recollected at the end of each session as a hormonal (progesterone and estradiol) control of the menstrual phase, but issues related to sample collection and storage hindered us from analyzing the samples.

3 Additionally, and in order to test whether the observed effects (or lack of effects) were due to recording or preprocessing strategies, an additional set of pre-processed data was prepared, whose details are described in <u>Supplementary Material</u>. It consisted in re-referencing electrodes offline to the common average, given that this type of reference may lead to greater N170 sensitivity to facial expressions (Hinojosa et al., 2015).

4 Also in this step, two additional quantification strategies were developed using the traditional average voltage quantification within windows-of-interest (WOI), provided that the PCA strategy may result in more conservative analytical outputs, according to our experience. Thus, this WOI plus direct voltage quantification was computed on i) nose tip-referenced data and ii) common average re-referenced data (see previous footnote). Details of both are described in <u>Supplemental Material</u>. Importantly, results derived from these alternative analyses supported those described in the main text on nose tip reference plus PCA analytical strategy.

5 <u>Supplementary Material</u> describes in detail the results derived from the alternative preprocessing and analytical strategies pointed out in footnotes 3 and 4. As may be appreciated, results regarding P1p, N170 and N2x from traditional quantification of both nose-tip and common-reference preprocessing confirm those described in the main text. As regards LPP, traditional analyses on nose-tip data do also confirm those described here, albeit those on common-reference data fail to find a significant effect of phase.

