## Supplementary Material for "Menstrual cycle and exogenous attention toward emotional expressions"

***Traditional average-voltage quantification of relevant ERP components, using the nose tip reference and average reference***

In order to guarantee that the observed effects, or lack of effects, were not due to pre-processing or analytical strategies, traditional voltage-based analyses were applied on both nose-tip- and average-referenced data.

#### Methods

After artifact removal, as explained in the Methods section, nose-tip-referenced EEG data were re-referenced to the average. Next, amplitudes of relevant ERP components were quantified through their mean voltage within a specified time window and topography. The time window and scalp region distinguishing each component were determined by visual inspection of grand averages and topographies of our data, following previous studies[1]. The mean voltages within each temporal window and at the relevant channels selected for each component were statistically analyzed through repeated-measures ANOVAs, considering Channel, Phase, and Emotion as factors. As in previous analyses, the levels of the factor Phase were FOL (the follicular phase) and LUT (the luteal phase), and the levels of the factor Emotion were Hap (happy faces), Ang (angry faces), and Neu (neutral faces). The levels of the factor Channels were the electrodes corresponding to the topography at which each component was quantified. Additionally, Bayesian repeated-measures ANOVAs were carried out on Phase × Emotion interactions for all ERP components, in order to obtain evidence in favor of the null hypothesis (Fol=Lut within all levels of Emotion) over the alternative hypothesis (Fol ≠ Lut within any level of Emotion). All statistical analyses were performed as explained in the Methods section.

[1] This traditional spatiotemporal characterization resulted similar to the one obtained by PCA.

### Results

1. <u>Experimental effects on ERP components using the nose tip reference</u>

Supplementary Table 1 shows means, standard deviations, and repeated-measures ANOVA results of P1p, N170, N2a, and LPP corresponding to each condition, when nose tip reference was used. Supplementary Figure 1 shows a selection of grand averages from each ERP.

| | Means (SD) | | | | | | Phase | | | Emotion | | | Phase x Emotion | | |
|---|---|---|---|---|---|---|---|---|---|---|---|---|---|---|---|
| | folH | folA | folN | lutH | lutA | lutN | F (1,42) | *p* | $\eta^2_p$ | F (2, 84) | *p* | $\eta^2 p$ | F (2,84) | *p* | $\eta^2_p$ |
| **P1p** | 3.88 (0.455) | 4.03 (0.438) | 3.81 (0.453) | 4.17 (0.557) | 4.19 (0.516) | 3.79 (0.505) | 0.287 | 0.595 | 0.007 | 1.834 | 0.166 | 0.042 | 0.391 | 0.678 | 0.009 |
| **N170 (right)** | -0.659 (0.576) | -0.454 (0.588) | -0.634 (0.520) | -0.618 (0.426) | -0.608 (0.432) | -0.641 (0.444) | 0.015 | 0.903 | 0 | 0.372 | 0.690 | 0.009 | 0.175 | 0.840 | 0.004 |
| **N170 (left)** | -0.561 (0.526) | -0.415 (0.568) | -0.790 (0.535) | -0.473 (0.500) | -0.677 (0.442) | -0.497 (0.441) | 0.019 | 0.891 | 0 | 0.3081 | 0.736 | 0.007 | 1.1543 | 0.320 | 0.027 |
| **N2a** | -3.74 (0.363) | -3.15 (0.353) | -3.37 (0.344) | -3.56 (0.304) | -3.39 (0.309) | -3.50 (0.348) | 0.039 | 0.844 | 0.001 | **3.943** | **0.023** | **0.086** | 1.176 | 0.314 | 0.027 |
| **LPP** | 4.18 (0.445) | 4.55 (0.501) | 4.60 (0.475) | 3.46 (0.566) | 3.78 (0.532) | 3.64 (0.456) | **9.761** | **0.003** | **0.189** | 2.301 | 0.106 | 0.052 | 0.197 | 0.822 | 0.005 |

**Supplementary Table 1.** Means and standard deviations (in parenthesis) of amplitudes for P1p, N170, N2a, and LPP corresponding to each condition when nose tip reference was used. Results of the repeated-measures ANOVAs with Phase and Emotion as factors. p values were GG-corrected when necessary. Significant and relevant results are shown in bold.

*P1p*: To analyze P1p, we established a temporal window from 120 to 145 ms. The channels selected according to the posterior topography of the P1p component were: PO7, PO3, PO1, POz, PO2, PO4, PO8, O1, Oz, and O2. Supplementary Figure 1 shows grand averages at POz highlighting the P1p temporal window. Repeated-measures ANOVAs on the main effect of Phase and Emotion, and the interaction effects showed no significant differences. Additionally, Bayesian repeated-measures ANOVA confirmed very strong evidence in favor of the null hypothesis regarding the Phase and Emotion interaction effect ($BF_{10}$=0.0266).

*N170*: To quantify this component, we established a temporal window from 160 to 200 ms. The channels selected according to the right distribution of the N170 component were: TP8, P8, and PO8. The channels selected to determine its left distribution were: TP7, CP5, and PO7. Figure 1 shows grand averages at P7 and P8, indicating the N170 temporal window. Repeated-measures ANOVAs on the main effect of Phase and the main effect of Emotion, as well as the interaction effects, revealed no significant differences either for the right or the left distribution of N170. Bayesian repeated-measures ANOVA confirmed strong evidence in favor of the null hypothesis regarding the Phase and Emotion interaction effect for the right ($BF_{10}$=0.0330) and the left ($BF_{10}$=0.0871) distribution.

*N2a*: To analyze this component, we selected a temporal window from 240 to 290 ms, choosing the following channels according to the anterior scalp region of the N2a component: Fp1, Fpz, Fp2, AF3, AFz, AF4, F3, F1, Fz, F2, and F4. Figure 1 shows grand averages at Fz and the N2a temporal window. Data analysis revealed no significant Phase × Emotion interaction, and Bayesian analysis regarding the Phase and Emotion interaction confirmed substantial evidence in favor of the null hypothesis ($BF_{10}$=0.12). No main effect of Phase was found. However, a significant effect of Emotion was observed [F (2, 84) = 3.942, $p$ = 0.023, $\eta^2_p$ =0.086]; post-hoc comparisons showed greater (more negative) amplitudes for Hap than Ang faces (t (42) = -3.03, Bonferroni corrected $p$=0.013).

*LPP*: To analyze LPP, we established a time window from 440 to 560 ms at P7, P5, P3, P1, Pz, P2, P4, P6, P8, PO7, PO3, PO1, POz, PO2, PO4, PO8, O1, Oz, and O2. Figure 1 shows grand averages at Pz and the selected window for LPP. This component showed no significant Phase × Emotion interaction, which was confirmed by Bayesian analysis showing decisive evidence in favor of the null hypothesis ($BF_{10}$=0.0058). A significant main effect of Phase [F (1, 42) = 9.761, $p$ = 0.003, $\eta^2_p$ = 0.189], but not of Emotion, was found. Post-hoc comparisons showed significantly greater LPP amplitudes in Fol than Lut (t(42) = 3.12, Bonferroni corrected $p$ = 0.003).

#### 2. Experimental effects on ERP components using the common average reference

Supplementary Table 2 shows means, standard deviations, and repeated-measures ANOVA results of P1p, N170, N2a, and LPP corresponding to each condition, when the common average reference was used. Supplementary Figure 2 shows a selection of grand averages from each ERP.

**Supplementary Table 2.** Means and standard deviations (in parenthesis) of amplitudes for P1p, N170, N2a, and LPP corresponding

| | **Means (SD)** | | | | | | **Phase** | | | **Emotion** | | | **Phase x Emotion** | | |
|---|---|---|---|---|---|---|---|---|---|---|---|---|---|---|---|
| | folH | folA | folN | lutH | lutA | lutN | F (1,42) | *p* | $\eta^2_p$ | F (2,84) | *p* | $\eta^2p$ | F (2,84) | *p* | $\eta^2_p$ |
| **P1p** | 4.23 (0.286) | 4.36 (0.283) | 4.18 (0.284) | 4.37 (0.366) | 4.27 (0.325) | 4.21 (0.337) | 0.024 | 0.877 | 0.001 | 1.219 | 0.301 | 0.028 | 1.171 | 0.315 | 0.027 |
| **N170 (right)** | -0.236 (0.367) | -0.033 (0.352) | -0.007 (0.343) | -0.180 (0.293) | -0.166 (0.285) | -0.007 (0.290) | 0.013 | 0.911 | 0 | 2.833 | 0.064 | 0.063 | 0.755 | 0.473 | 0.018 |
| **N170 (left)** | -0.149 (0.280) | -0.078 (0.298) | -0.286 (0.277) | -0.067 (0.290) | -0.161 (0.265) | -0.167 (0.271) | 1.336 | 0.254 | 0.031 | 0.413 | 0.663 | 0.010 | 6.154 | 0.003 | 0.128 |
| **N2a** | -2.49 (0.289) | -2.38 (0.274) | -2.56 (0.270) | -2.34 (0.268) | -2.10 (0.248) | -2.44 (0.248) | 0.907 | 0.346 | 0.021 | **8.123** | **<0.001** | **0.162** | 0.695 | 0.502 | 0.016 |
| **LPP** | 1.168 (0.247) | 1.435 (0.262) | 1.354 (0.263) | 0.998 (0.222) | 1.097 (0.222) | 1.069 (0.212) | 1.76 | 0.191 | 0.040 | **4.59** | **0.013** | **0.098** | 1.12 | 0.331 | 0.026 |

to each condition, when the common average reference was used. Results of repeated-measures ANOVAs with Phase and Emotion as factors. p values were GG-corrected when necessary. Significant and relevant results are shown in bold.

*P1p*: To quantify P1p, we established a time window from 120 to 145 ms. The channels we selected according to the posterior distribution of the P1p component were: PO7, PO3, PO1, POz, PO2, PO4, PO8, O1, Oz, and O2. Figure 2 shows the grand averages at POz and the window in which P1p was scored. Repeated-measures ANOVAs on the main effect of Phase, the main effect of Emotion, and the interaction effects yielded no significant differences. Additionally, Bayesian repeated-measures ANOVA confirmed decisive evidence in favor of the null hypothesis regarding the Phase and Emotion interaction effect ($BF_{10}$<0.01).

*N170*: To analyze this component, we established a temporal window from 160 to 200 ms. The channels selected to determine the right topography of the N170 component were TP8, P8, and PO8, and for the left distribution TP7, CP5, and PO7. Supplementary Figure 2 shows grand averages at P7 and P8, also indicating the N170 temporal window. Repeated-measures ANOVAs on the main effect of phase, the main effect of emotion, and the interaction effects found no significant differences for the right distribution of N170. In addition, Bayesian repeated-measures ANOVA confirmed strong evidence in favor of the null hypothesis regarding the Phase × Emotion interaction effect ($BF_{10}$=0.0333). The left distribution showed a significant effect of Phase × Emotion [$F(2, 84) = 6.154$, $p = 0.003$, $\eta^2_p = 0.128$], but post hoc tests failed to reach significance [$p > 0.05$ in all cases]. Likewise, Bayesian ANOVA regarding the phase and emotion interaction found substantial evidence in favor of the null hypothesis ($BF_{10}$=0.133). No main effect of Phase or Emotion was found. A significant effect of Channel × Emotion [$F(4, 168) = 3.963$, G-G corrected $p = 0.009$, $\eta^2_p = 0.086$, Epsilon=0.826] was also found, although post hoc tests again failed to reach significance for all the relevant comparisons [$p > 0.05$ in all cases].

*N2a*: To analyze this component, we established a temporal window from 230 to 280 ms. The channels selected according to the anterior distribution of the N2a component were: Fp1, Fpz, Fp2, AF3, AFz, AF4, F3, F1, Fz, F2, and F4. Figure 2 shows grand averages at Fz and the selected temporal window. Traditional ANOVA analysis revealed no significant Phase × Emotion interaction. Bayesian analysis regarding the Phase and Emotion interaction confirmed decisive evidence in favor of the null hypothesis ($BF_{10}$=0.0051). No main effect of Phase was found. However, a significant effect of Emotion was observed [$F(2, 84) = 8.123$, $p < 0.01$, $\eta^2_p$ =0.162]. Post-hoc comparisons showed significant differences consisting in greater (i.e., more negative) amplitudes in case of Hap and Neu, compared to Ang faces ($t(84) = -2.94$, Bonferroni corrected $p$=0.016 and $t(88) = 3.85$, Bonferroni corrected $p$=0.001, respectively).

*LPP*: To quantify the LPP component, we fixed a temporal window from 440 to 560 ms at a posterior topography including P7, P5, P3, P1, Pz, P2, P4, P6, P8, PO7, PO3, PO1, POz, PO2, PO4, PO8, O1, Oz, and O2. Supplementary Figure 2 shows grand averages at Pz, highlighting the chosen temporal window. This component showed no Phase × Emotion interaction in repeated-measures ANOVA. In addition, Bayesian analysis on this interaction revealed decisive evidence in favor of the null hypothesis ($BF_{10}$=0.0024). A significant main effect of Emotion [$F(1, 42) = 4.59$, $p = 0.014$, $\eta^2_p = 0.098$], but not of Phase, was found. Post-hoc comparisons showed significantly greater LPP amplitudes in Ang than Hap ($t(42) = 3.145$, Bonferroni corrected $p = 0.009$).

#### Discussion and conclusion

These traditional voltage-based analyses were performed in order to replicate the results obtained using Principal Component Analysis (PCA) on nose-tip-referenced EEG data, given that i) traditional voltage-based quantification may yield less conservative outputs than PCA, and ii) the common average reference may provide better sensitivity of the N170 component to facial expressions than the nose tip reference. As a result, ANOVAs did not reveal any significant modulation of menstrual phase on exogenous attention to emotional expressions for both nose tip and common average-referenced data, in line with our previous PCA results. Additionally, Bayesian analysis provided evidence in favor of the null hypothesis, confirming the absence of modulation in all relevant ERP components.

Moreover, concerning our secondary results, the effect of emotion on N2a was also confirmed when employing traditional analyses and for both references. On the contrary, in the case of LPP, we obtained the same results showing a modulation of Phase, when using the nose tip reference, but not with the average reference, where a main effect of Emotion was evident.

Modulations by the emotional content of distractors on LPP -though not being the most common result- have repeatedly been described in literature, in particular showing an advantage of negative stimuli (Nordström & Wiens, 2012; Schönwald & Müller, 2014; Wiens et al., 2011). In sum, we cannot discard certain variability underlying our LPP results when modifying pre-processing and analytical strategies, thus, these secondary control results should be taken with caution.

However, outcomes concerning our main hypothesis considering the absence of modulation of exogenous attention by the menstrual cycle in the components sensitive to our experimental task (P1p, N170, and the N2x family) seem to be independent of decisions on pre-processing and analysis. Hence, these new analyses confirm the absence of an interaction effect between phase and emotion, thus, confirming our hypothesis.

***References***


Nordström, H., & Wiens, S. (2012). Emotional event-related potentials are larger to figures than scenes but are similarly reduced by inattention. BMC neuroscience, 13(1), 1-10.

Schönwald, L. I., & Müller, M. M. (2014). Slow biasing of processing resources in early visual cortex is preceded by emotional cue extraction in emotion–attention competition. Human Brain Mapping, 35(4), 1477-1490.

Wiens, S., Sand, A., Norberg, J., & Andersson, P. (2011). Emotional event-related potentials are reduced if negative pictures presented at fixation are unattended. Neuroscience letters, 495(3), 178-182.

#### Supplementary Figures

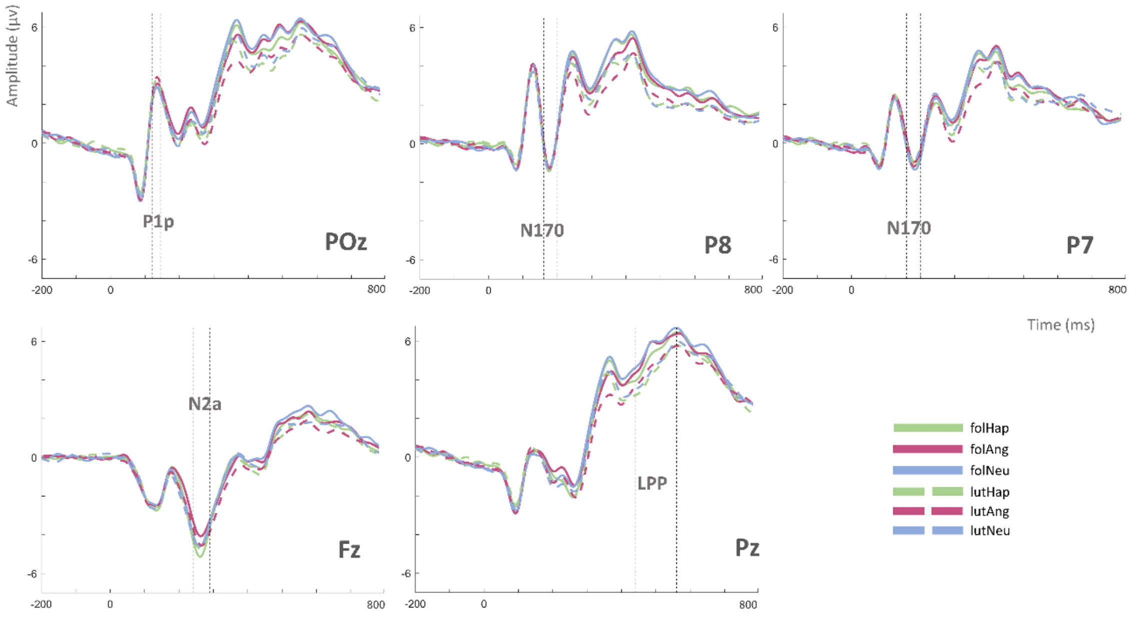


**Supplementary Figure 1**. Grand averages corresponding to posterior (POz and Pz), anterior (Fz), and temporo-parietal (P7 and P8) electrodes, including the temporal windows for P1p, N170, N2a, and LPP, when nose tip reference was used.

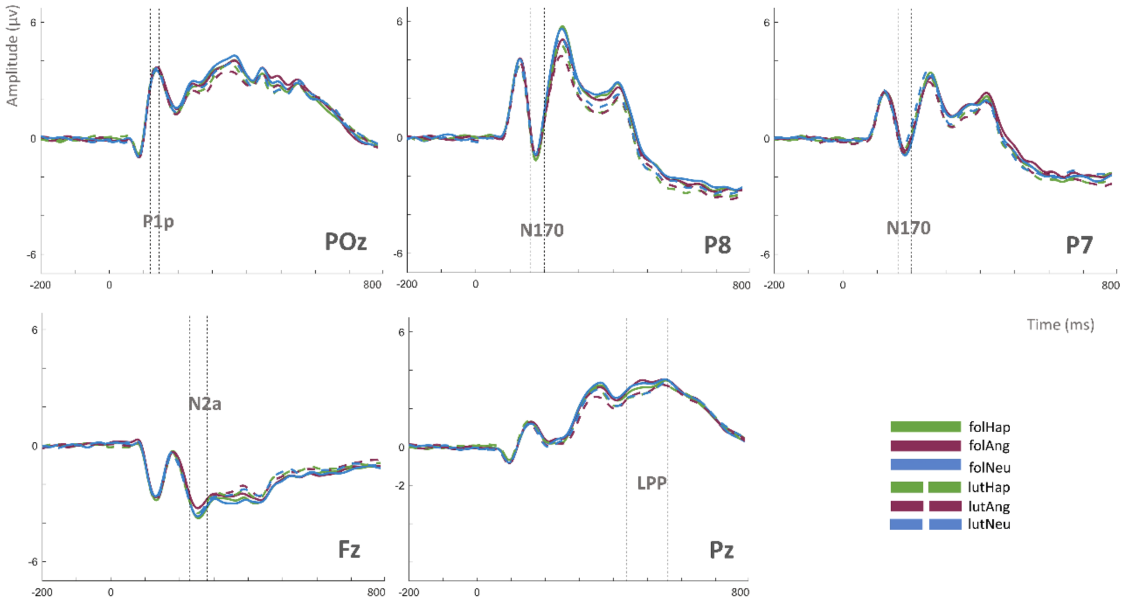


**Supplementary Figure 2**. Grand averages corresponding to posterior (POz and Pz), anterior (Fz), and temporo-parietal (P7 and P8) electrodes, highlighting the temporal windows for P1p, N170, N2a, and LPP, when common average reference was sed.
